# Can hot temperatures limit disease transmission? A test of mechanisms in a zooplankton-fungus system

**DOI:** 10.1101/605477

**Authors:** Marta S. Shocket, Alexandra Magnante, Meghan A. Duffy, Carla E. Cáceres, Spencer R. Hall

**Author notes:** Corresponding Author: Marta S. Shocket. Department of Ecology and Evolutionary Biology, University of California, Los Angeles, CA, USA. **Author Contributions**: SRH, CEC, and MAD designed and obtained funding for the field survey. SRH and MSS collected field data. MSS and SRH designed the laboratory studies; MSS and AM conducted them. MSS wrote the first draft of the manuscript, and all authors revised and approved the final version. **Data Accessibility**: Upon acceptance all data and code will be stored in Dryad Data Repository.

## Abstract

1. unimodally to temperature, i.e., be maximized at intermediate temperatures and constrained at extreme low and high temperatures. However, empirical evidence linking hot temperatures to decreased transmission in nature remains limited.
2. We tested the hypothesis that hot temperatures constrain transmission in a zooplankton-fungus (*Daphnia dentifera-Metschnikowia bicuspidata*) disease system where autumnal epidemics typically start after lakes cool from their peak summer temperatures. This pattern suggested that maximally hot summer temperatures could be inhibiting disease spread.
3. Using a series of lab experiments, we examined the effects of high temperatures on five mechanistic components of transmission. We found that (1) high temperatures increased exposure to parasites by speeding up foraging rate but (2) did not alter infection success post-exposure. (3) High temperatures lowered parasite production (due to faster host death and an inferred delay in parasite growth). (4) Parasites made in hot conditions were less infectious to the next host (instilling a parasite ‘rearing’ or ‘trans-host’ effect of temperature during the prior infection). (5) High temperatures in the free-living stage also reduce parasite infectivity, either by killing or harming parasites.
4. We then assembled the five mechanisms into an index of disease spread. The resulting unimodal thermal response was most strongly driven by the rearing effect. Transmission peaked at intermediate-hot temperatures (25-26°C) and then decreased at maximally hot temperatures (30-32°C). However, transmission at these maximally hot temperatures only trended slightly lower than the baseline control (20°C), which easily sustains epidemics in laboratory conditions and in nature. Overall, we conclude that while exposure to hot epilimnetic temperatures does somewhat constrain disease, we lack evidence that this effect fully explains the lack of summer epidemics in this natural system. This work demonstrates the importance of experimentally testing hypothesized mechanisms of thermal constraints on disease transmission. Furthermore, it cautions against drawing conclusions based on field patterns and theory alone.

## INTRODUCTION

How do high temperatures affect the spread of infectious diseases? In the current prevailing view, warming from climate change will shift the geographic range of diseases: some new areas will become warm enough to support disease, whereas others that previously sustained disease will become too hot (Altizer, Ostfeld, Johnson, Kutz, & Harvell, 2013; Lafferty, 2009; Lafferty & Mordecai, 2016). This hypothesis stems from a principle of thermal biology: most biological traits have unimodal reaction norms, where performance peaks at intermediate temperatures and declines to zero at cooler and warmer temperatures (Dell, Pawar, & Savage, 2011). Thus, once temperatures exceed the thermal optima of traits driving transmission, disease should decline. Many models predict upper thermal constraints on diseases, e.g., helminthic ungulate parasites (Molnár, Kutz, Hoar, & Dobson, 2013), a rhizocephalan crab parasite (Gehman, Hall, & Byers, 2018), a microsporidian *Daphnia* parasite (Kirk et al., 2018), schistosomiasis (Mangal, Paterson, & Fenton, 2008), and mosquito-borne pathogens (Mordecai et al., *in press*). Additionally, there is evidence for upper thermal constraints on disease in natural populations of the crab parasite (Gehman et al., 2018), mosquito-borne pathogens (e.g., Shocket et al., *in review*) and fungi infecting grasshoppers (Carruthers, Larkin, & Firstencel, 1992), amphibians (Berger et al., 2004; Raffel, Michel, Sites, & Rohr, 2010), and bats (Langwig et al., 2015). However, temperature often co-varies with other seasonal environmental factors, so causally linking temperature to observed patterns of disease is challenging (Altizer et al., 2006; Pascual & Dobson, 2005). Thus, the generality of upper thermal constraints excluding disease remains unclear.

Conceptually, upper thermal constraints act like fever, taking advantage of a common thermal mismatch between hosts and parasites. Because hosts can often endure hotter environments than their parasites, many animals increase their body temperature when infected (see citations below). In ectotherms, fever arises from behavioral thermoregulation (microhabitat selection) and is widespread, occurring in vertebrates (including amphibians, reptiles, and fish: Rakus, Ronsmans, & Vanderplasschen, 2017), snails (Zbikowska, Wrotek, Cichy, & Kozak, 2013), and insects (including bees, flies, grasshoppers, mosquitoes, and beetles: Stahlschmidt & Adamo, 2013; Thomas & Blanford, 2003). Behavioral fever can impair parasite performance, enhancing clearance or reducing virulence of infection. An analogous process can occur within ectothermic hosts inhabiting high ambient temperatures (regardless of infection status)—in essence, an environmental fever. High ambient temperatures can also harm parasites with free-living stages outside of hosts. Mechanistically linking high temperatures to reduced disease requires examining thermal effects on components of the transmission process (McCallum et al. 2017). We use the term ‘transmission (process)’ to broadly refer to the full parasite life cycle, including infective propagule production and propagule survival in the environment; we also use ‘transmission rate’ narrowly defined as the rate of new infections (i.e., the parameter ‘*β’* calculated from infection prevalence and densities of hosts and parasites; McCallum et al. 2017).

Here, we use a series of experiments to evaluate mechanisms for potential upper thermal constraints on transmission in a planktonic-fungal disease system. Autumnal epidemics start once lake waters cool below summer maxima (Fig. 1A). These delayed starts could reflect hot temperatures inhibiting disease if they push any of five transmission components past their thermal optima (Fig. 1B). First, hot temperatures could slow host feeding and lower consumption-based exposure to parasites. Second, hot temperatures could reduce parasite infectivity inside hosts, lowering the probability of successful infection (via effects on hosts and/or parasites). Third, hot temperatures could decrease the quantity of parasite propagules [spores] produced by an infection. This decrease could stem from slower host growth rate (since parasite production often scales with host growth: Hall, Knight, et al., 2009; Hall, Simonis, Nisbet, Tessier, & Cáceres, 2009), slower parasite growth independent from host growth, or enhanced mortality of infected hosts (truncating production time; Auld, Hall, Housley Ochs, Sebastian, & Duffy, 2014; Civitello, Forys, Johnson, & Hall, 2012). Fourth, hot temperatures could lower the quality of parasite spores released from dead hosts into the environment (Shocket, Vergara, et al., 2018). Finally, these free-living spores could be harmed or killed by hot temperatures. Thus, high temperatures could constrain this fungal disease at multiple stages of the transmission process.

**Figure 1:**
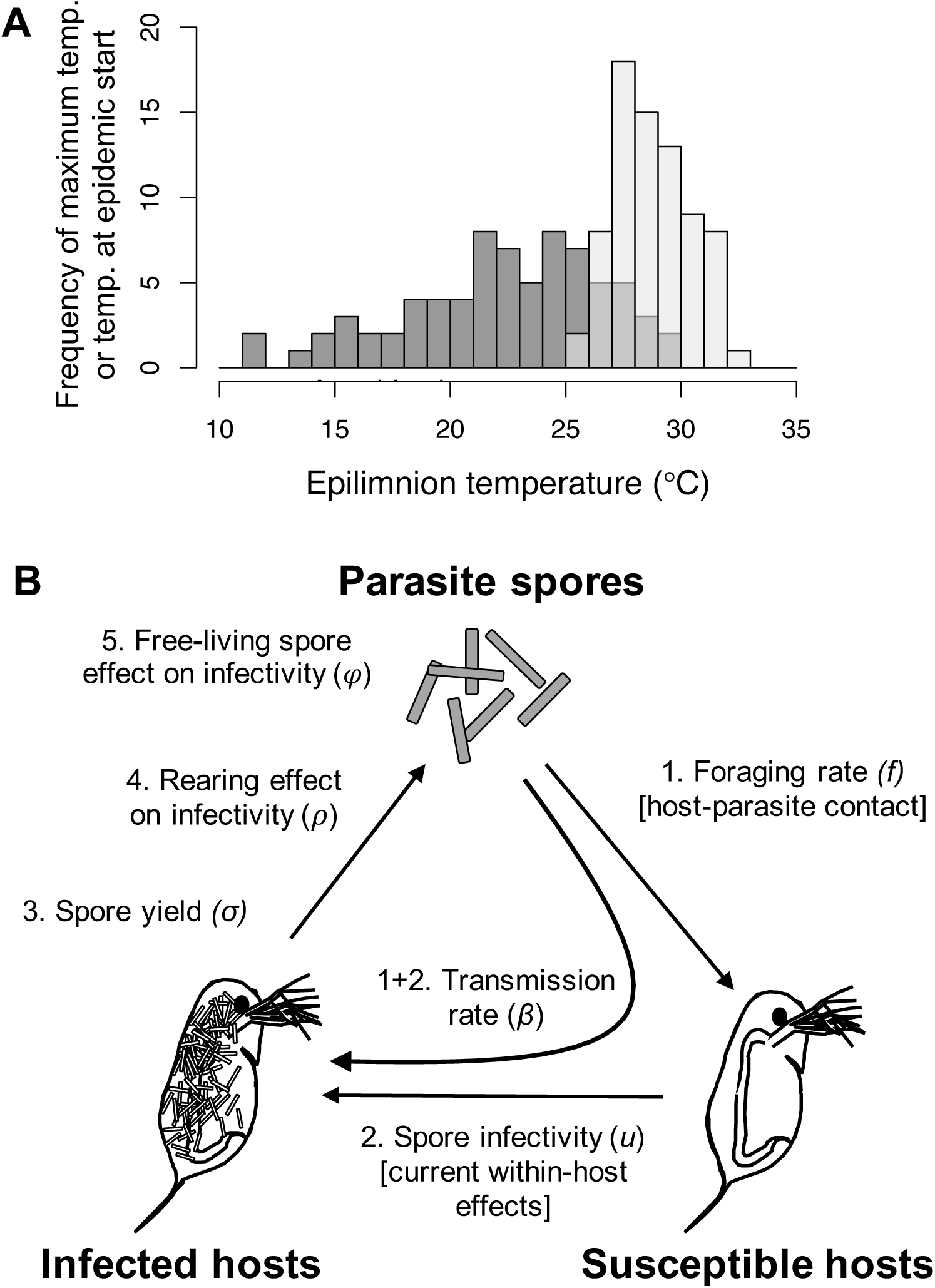
Motivating field pattern and mechanistic components of transmission. (A) Fungal epidemics usually start (dark grey bars) after lakes have cooled from the maximum summer temperature (light grey bars). Epidemics never started when the epilimnion (upper, warmer layer) was hotter than 30°C, suggesting an upper thermal constraint. Data summarize 74 epidemics from 20 lakes in Indiana (USA) sampled from 2009-2015. (B) High temperature could limit transmission via five mechanisms. 1-2) Hosts become infected at transmission rate *β*, which can be divided into 1) host foraging rate (*f*), i.e., exposure to spores, and 2) spore infectivity, as determined by within-host processes (*u*). 3) Parasite spores are produced at spore yield (*σ*). 4) A rearing effect from temperature during the previous infection (ρ) determines initial spore infectivity. 5) Harm to free-living spores (φ) might also impact their infectivity. The product of all five components (*f u σ ρ φ*) determines ‘transmission potential’.

## STUDY SYSTEM

The hosts (*Daphnia dentifera*) are zooplankton grazers in freshwater temperate lakes across the Midwestern United States; the fungal parasite *Metschnikowia biscupidata* causes epidemics in some host populations, with prevalence reaching up to 60% (Penczykowski, Hall, Civitello, & Duffy, 2014). Hosts become infected when they filter-feed on algae and inadvertently consume fungal spores (Hall et al., 2007). The spores pierce the host’s gut wall, entering its body cavity. Inside, fungal conidia replicate in the hemolymph before maturing into new spores (Stewart Merrill & Cáceres, 2018). Following host death, spores are released into the water for new hosts to consume (Ebert, 2005).

The seasonality of epidemics motivated a focus on high temperatures. Epidemics typically begin in late summer or early fall (August–October) and wane in late fall or early winter (November–December; Penczykowski, Hall, et al., 2014). During this time, lake water temperature declines (Shocket, Strauss, et al., 2018). Many traits that influence disease spread (host demographic traits, transmission rate, and spore production) change plastically with temperature (Hall, Tessier, Duffy, Huebner, & Cáceres, 2006; Shocket, Strauss, et al., 2018). Transmission increases with constant temperatures up to 26°C, and hosts cannot be cultured in constant temperatures above 27°C (Shocket, Strauss, et al., 2018). However, organisms can withstand otherwise lethal temperatures in fluctuating environments (Niehaus, Angilletta, Sears, Franklin, & Wilson, 2012). For instance, in our stratified lakes, hosts experience temperatures exceeding 27°C in summer (typical maxima 29–32°C; Fig. 1A): they migrate between the colder, deeper hypolimnion during day (to avoid mortality from visually-oriented fish predators) and the warmer, upper epilimnion at night (to take advantage of greater algal resources and faster growth in warmer temperatures) (Hall, Duffy, Tessier, & Cáceres, 2005; Lampert, 1989). Epidemics often begin as lakes start cooling from maximum summer temperatures (Fig. 1A). This pattern suggested that high temperatures could constrain disease spread, as predicted by theory (Lafferty, 2009; Lafferty & Mordecai, 2016).

## METHODS

### Field Survey

Field survey data generated the motivating pattern (Fig. 1A: the relationship between epidemic start date and epilimnetic temperature). We surveyed 10–28 lakes in Indiana (Greene and Sullivan Counties) weekly (2009–2011) or bi-weekly (2013–2014) from August to December. For each visit, we collected a zooplankton sample (13 cm diameter net with 153 µm mesh) and measured lake water temperature data at 0.5–1 meter intervals with a Hydrolab multiprobe (Hach Environmental). For each sample, we visually diagnosed 400+ live hosts with a dissecting scope (20-50X magnification). An epidemic ‘started’ when infection prevalence first exceeded 1% for two consecutive sampling visits (Shocket, Strauss, et al., 2018). We calculated the epilimnetic temperature by fitting a spline to temperature across water depth, and averaging from the water surface to the depth where the temperature gradient first exceeded 1°C m^−1^ (i.e., the thermocline; see Hite et al. 2016 Appendix S2).

### General Approach

We measured how high temperatures influence five components of the transmission process with laboratory assays (Table 1). Then we combined them into a synthetic index of disease spread: ‘transmission potential’ (Auld et al., 2014). For mechanisms involving the host or host-parasite interaction (mechanisms 1-3: foraging rate [*f*], spore infectivity from within-host processes [*u*], and spore yield [*σ*]), we used fluctuating temperatures to expose hosts to high temperatures for part of the day (they cannot survive constant temperatures >27°C). Hosts were kept on a 16:8 light:dark cycle. All hosts experienced the same 20°C temperature for 8 hours, and then 20, 26, or 32°C for 16 hours (‘maximum temperature’). For mechanisms 4-5 (rearing effect on spore quality [*ρ*] and free-living spore effect [*φ*]), we conducted common garden infection assays, exposing uniform hosts at constant 20°C to spores from different treatments. Thus, variation in transmission rate can be attributed to differences in spore infectivity. Temperatures varied slightly among experiments (25 or 26°C, 30 or 32°C) based on incubator availability. For calculating transmission potential, we treat temperature categorically and pool these treatments.

**Table 1:**
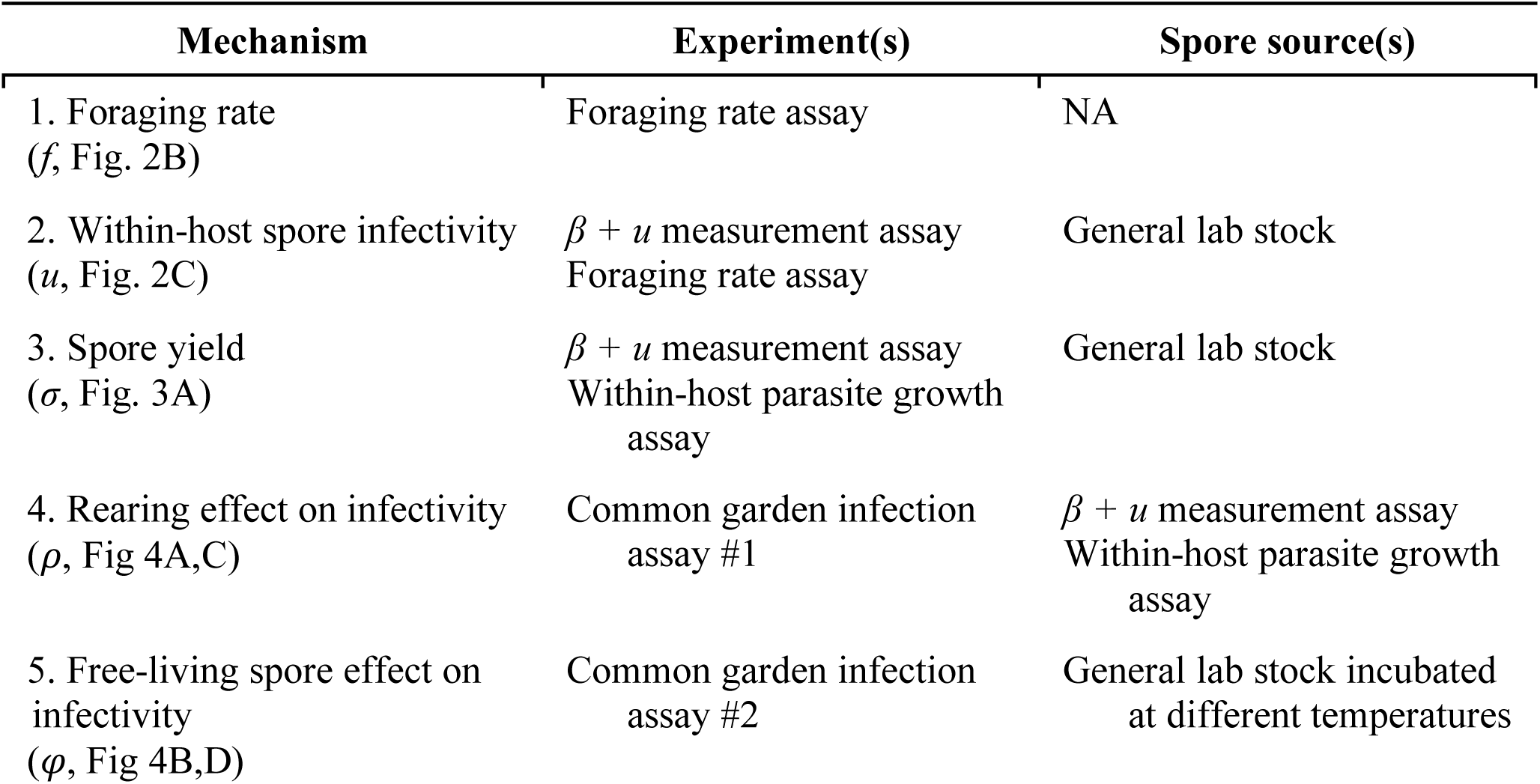
The experiments (and spore sources) used to test the five mechanistic components of disease transmission.

| Mechanism | Experiment(s) | Spore source(s) |
| --- | --- | --- |
| 1. Foraging rate<br>($f$, Fig. 2B) | Foraging rate assay | NA |
| 2. Within-host spore infectivity<br>($u$, Fig. 2C) | $\beta + u$ measurement assay<br>Foraging rate assay | General lab stock |
| 3. Spore yield<br>($\sigma$, Fig. 3A) | $\beta + u$ measurement assay<br>Within-host parasite growth assay | General lab stock |
| 4. Rearing effect on infectivity<br>($\rho$, Fig 4A,C) | Common garden infection assay #1 | $\beta + u$ measurement assay<br>Within-host parasite growth assay |
| 5. Free-living spore effect on infectivity<br>($\varphi$, Fig 4B,D) | Common garden infection assay #2 | General lab stock incubated at different temperatures |

Due to time and incubator constraints, we were unable to replicate experiments across multiple incubators. Thus, our temperature treatments are ‘pseudo-replicated’ in that all replicates for a treatment were conducted in the same incubator at the same time. Accordingly, our results may be influenced by random incubator effects.

### Mechanisms 1 & 2: Foraging rate (*f*) and spore infectivity from within-host processes (*u*)

We measured foraging rate of hosts by comparing the fluorescence of ungrazed and grazed algae (Penczykowski, Lemanski, et al., 2014; Sarnelle & Wilson, 2008). We added estimates of foraging rate at 30°C to those at 20 and 25°C presented elsewhere using the same methods (Shocket, Vergara, et al., 2018). In both experiments, we measured foraging rate across a gradient of host body size (Kooijman, 2009) to index foraging at a common size among experiments (1.5 mm). We used maximum likelihood estimation (MLE) to fit size-dependent functions of foraging with the ‘bbmle’ package (Bolker & R Development Core Team, 2017) in R (R Core Team, 2017). See Appendix for details.

We measured how high temperature impacts transmission rate (*β*) and spore infectivity from within-host processes (*u*) with an infection assay (‘*β + u* measurement assay’). For successful infection, the fungus must break through the host gut barrier and then replicate and develop within the host hemolymph. High temperatures could inhibit the parasite during either process. Thus, we factorially manipulated the maximum temperature (20 and 32°C) during parasite exposure and infection establishment (for four exposure/establishment treatments: 20/20, 20/32, 32/20, and 32/32°C) to reveal if high temperatures interfere at either step (similar to Allen & Little, 2011). Hosts were exposed individually in their *‘*exposure temperature*’* for 24 hours, then moved to their ‘infection establishment temperature.’ Later, hosts were visually diagnosed for infection. Transmission rate was calculated from proportion infected (see Appendix). We calculated spore infectivity from within-host processes (*u*) for each treatment by dividing transmission rate (*β*) by foraging rate (*f*) at the exposure temperature (*u*=*β*/*f*).

### Mechanism 3: Spore yield (*σ*) and related host and parasite traits

We measured how high temperatures impact final spore yield (*σ*) of infected hosts that died from their infection. This trait estimates spore release into the environment. We pooled spore yields from the *β + u* measurement assay (above; treatments = 20/20 and 32/32°C) and the within-host parasite growth assay (below; treatments = 20, 26, and 32°C) since they did not differ statistically (20°C: *p*=0.65; 32°C *p*=0.93). We tested for differences between temperatures by fitting a suite of models via MLE: in each model spore yield was normally distributed and temperature treatments could exhibit the same or different means and standard deviations. We compared models using AIC and calculated *p*-values with likelihood ratio tests.

To distinguish between three possible mechanisms driving the thermal response of spore yield, we quantified related host and parasite traits. First, we measured host growth rate (*g*_*h*_) with a juvenile growth rate assay (Lampert & Trubetskova, 1996)(see Appendix), since spore yield often scales with *g*_*h*_ (e.g., with different host food resources: Hall, Knight, et al., 2009; Hall, Simonis, et al., 2009). We compared treatments with t-tests. Second, we measured parasite growth (i.e., number of mature spores within hosts over time) using a sacrifice series (‘within-host parasite growth assay;’ see Appendix), since spore yield could decline if the number of parasites increases more slowly, independently of host condition (Thomas & Blanford, 2003). We fit and bootstrapped linear models of ‘spore load’ over time to estimate parasite growth rate (*g*_*p*_, the model slope). ‘Spore load’ estimates included spores in living (i.e., sacrificed) hosts, unlike ‘spore yield,’ which was calculated only from dead hosts that were killed by the parasite. Spore yield is directly relevant for the epidemiology of the system, while spore load measures an underlying process (parasite growth rate per day, *g*_*p*_) that contributes to spore yield. Spore load increased linearly over the full time series at 26 and 32 ° C. Spore load plateaued after day 19 at 20°C, so we truncated the time series to estimate the linear slope for only that portion. Finally, we calculated death rate (*d*) of infected hosts (see Appendix), since spore yield can decline with shorter host lifespan (Auld et al., 2014; Civitello et al., 2012). We compared treatments with randomization tests.

### Mechanisms 4 & 5: rearing (*ρ*) and free-living spore (*φ*) effects on infectivity

We measured how high temperatures modify spore infectivity prior to encountering hosts via a rearing effect on baseline spore quality (*ρ*) and harm to free-living spores (*φ*). We conducted infection assays on ‘common garden’ groups of hosts at 20°C using different spore treatments (i.e., from different spore rearing temperatures for *ρ* and from different spore incubation temperatures for *φ*). Thus, variation in transmission rate reflects differences in spore infectivity. To measure *ρ*, we conducted two experiments, one with spores produced in the *β + u* measurement assay (20/20 and 32/32°C treatments) and another with spores produced in the within-host parasite growth assay (20, 26, and 32°C treatments). To measure *φ*, we used spores incubated at three temperatures (20, 25, and 30°C) for two durations (1-day and 7-days) in constant, non-fluctuating temperatures (spores do not migrate between stratified water layers). One-day incubations were stored at 4°C for the first six days (standard procedure for spore storage). We estimated transmission rates (*β*) from the prevalence data (see Appendix).

Both mechanisms influence transmission by modifying spore infectivity (already estimated from within-host processes as *u*, mechanism 1). Thus, in order to incorporate these mechanisms into a synthetic metric for disease spread (transmission potential, see below), we calculated unit-less rearing (*ρ*) and free-living (*φ*) effects standardized to infectivity at 20°C. Specifically, we calculated the parameters by dividing the estimates for transmission rate (*β*) at 26 and 32°C by that at 20°C. Accordingly, values of *ρ*<1 or *φ*<1 mean spores are less infectious due to rearing or free-living effects than at 20°C, respectively; conversely, values >1 mean spores are more infectious than at 20°C. To calculate confidence intervals at 20°C, we divided a bootstrapped distribution of transmission rates by a randomly-shuffled version of itself. Additionally, because harm to free-living spores occurs over time as spores are removed by hosts, we used a simple model to estimate time-weighted transmission rates for *φ*. We assumed that spore infectivity declined linearly over the 7-day assay, and that hosts consume spores at a constant foraging rate (resulting in an exponential decay in spores remaining over time). Thus, we weighted the estimated transmission rate on each day by the proportion of spores consumed by hosts on that day (see Appendix for detailed methods and a sensitivity analysis of the model).

### Transmission potential

We calculated an index of disease spread to synthesize the effects of all five mechanisms. We defined transmission potential as the product of all five parameters (*f u σ ρ φ*). We generated confidence intervals using bootstrapped parameter distributions. To visualize the contribution of each parameter, we calculated transmission potential for each of the five possible four-parameter combinations, holding the fifth parameter constant at its 20°C point estimate. These values reveal how each parameter affects the magnitude and uncertainty of transmission potential (i.e., a type of sensitivity analysis).

### Additional Statistical Analyses

For all parameters, we bootstrapped 95% confidence intervals (data sampled within groups, with replacement; 10,000 samples). For parameters derived from transmission rates (*β, u, ρ*, and *φ*), we used randomization tests to compare temperature treatments, since a single value is calculated from all individuals (treatment labels shuffled among host individuals, without replacement; 10,000 samples). For *f* and transmission potential (for which traditional statistical tests were not available), we used the bootstrapped distributions to compare treatments. Specifically, we calculated the cumulative probability density of the best estimate from one treatment according to the bootstrapped distribution of the other. These ‘PD-values’ are analogous to *p*-values. We considered treatments significantly different if *PD*<0.025. See Appendix for details and a complete list of statistical tests and results.

## RESULTS

### Mechanisms 1 & 2: Foraging rate (*f*) and spore infectivity from within-host processes (*u*)

Contrary to our predictions, high temperature did not lower transmission rate (*β*) during either step (exposure or infection establishment; Fig. 2A). Instead, transmission rate was higher when hosts were exposed at 32°C than at 20°C (20°C infection establishment: *p*=0.0013; 32°C infection establishment: *p*<0.0001). Temperature during infection establishment exerted no effect on transmission rate (20°C exposure: *p*=0.10; 32°C exposure: *p*=0.31). When exposure and establishment temperatures were equal (as in nature; the 20/20 and 32/32°C treatments here), transmission rate was higher at 32°C than at 20°C (*p*=0.0068). Thus, even at maximal epilimnetic temperatures, the impacts of higher temperatures on transmission rate promoted rather than inhibited disease.

**Figure 2:**
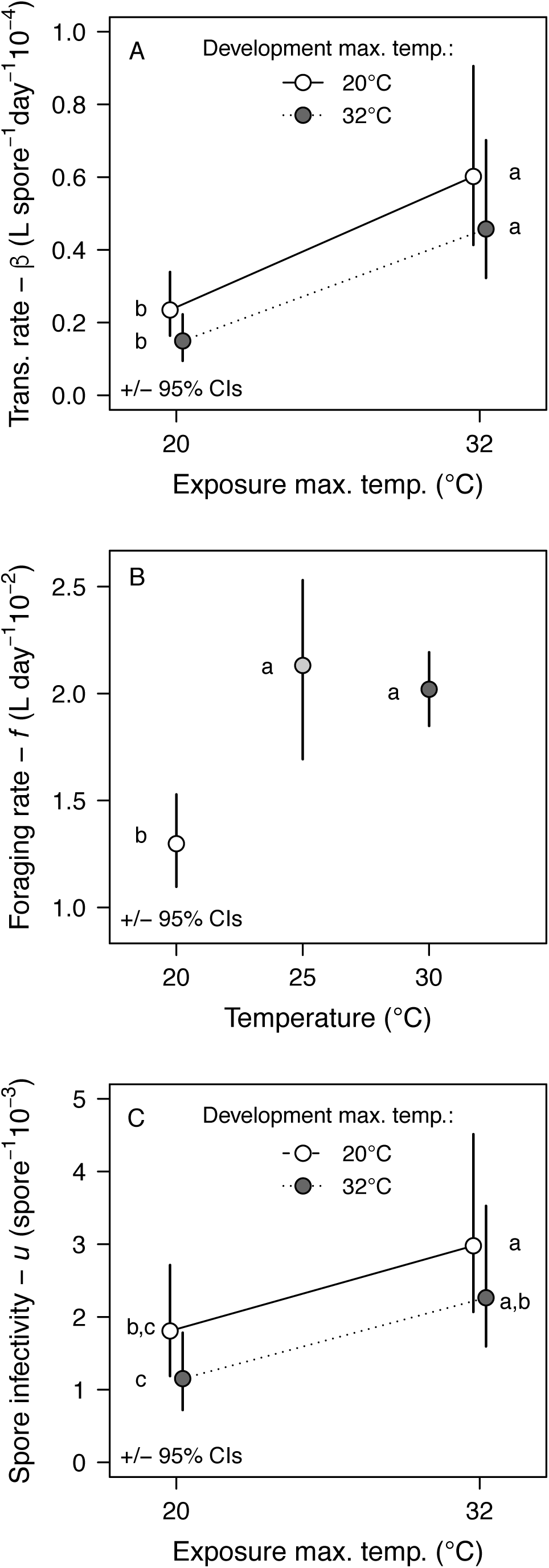
High temperature impacts on transmission rate (*β*), foraging rate (*f*, mechanism 1) and spore infectivity from current within-host processes (*u*, mechanism 2). In A and C, the effect of high temperature during parasite exposure and infection establishment (20°C infection establishment = white circles, solid line; 32°C infection establishment = dark grey circles, dotted line). (A) Transmission rate (*β*) increased when hosts were exposed at 32°C and did not change with infection establishment temperature. For constant temperatures, transmission is higher at 32°C than at 20°C. (B) Foraging (exposure) rate of hosts (*f*) is higher at 26°C (light grey) and 32°C (dark grey) than at 20°C (white). (C) Spore infectivity (*u*=*β*/*f*) increased when hosts were exposed at 32°C for both infection establishment temperatures. However, for constant temperatures, infectivity did not differ between 20 and 32°C. Error bars show 95% CIs. Letters indicate significant differences.

The thermal response of transmission rate was mechanistically driven by foraging rate of hosts (*f*), not spore infectivity from within-host processes (*u*). Foraging rate increased from 20 to 25°C (*PD*=0; see *Methods* and Appendix for a description of PD values, which are analogous but not identical to *p*-values) and then plateaued at 30°C (*PD*=0.11; Fig. 2B). Thus, hosts encounter more spores at 25 and 30°C than at 20°C. After we accounted for predicted host-parasite contact, spore infectivity was fairly insensitive to high temperatures (Fig. 2C). Temperature during infection establishment did not impact spore infectivity (20°C exposure: *p*=0.10; 32°C exposure: *p*=0.31). Exposure temperature increased spore infectivity (20°C infection establishment: *p*=0.034; 32°C infection establishment: *p*=0.0052), but in the opposite direction of the hypothesized mechanism (hotter temperature increased infectivity). When exposure and infection establishment temperatures were equal (as in nature), spore infectivity did not differ (*p*=0.37). Thus, high temperatures increased the foraging rate of hosts, elevating host contact with spores, while spore infectivity barely changed. These changes in parasite exposure led to more transmission at high temperatures.

### Mechanism 3: Spore yield (*σ*) and other measures of host and parasite growth

Final spore yield (*σ*) in hosts that died from infection was lower at 32°C than at 20 and 26°C (Fig. 3A; best-fitting model had two means, see Table S5 for model AIC scores and Akaike weights). This pattern was not explained by host condition estimated via growth rate. Host growth rate (*g*_*h*_, Fig. 3B) always increased with temperature (20 versus 26°C: *p*=4.7 × 10^−6^; 26 vs. 32°C: *p*=0.00038). Instead, the pattern was explained by a combination of host death rates and delays in spore maturation. Infected hosts died more quickly at 26°C than 20°C (*p*<0.0001), and death rate trended higher from 26 to 32°C (*p*=0.063; Fig. 3C). Meanwhile, growth rate of mature parasite spores (*g*_*p*_, time series in Fig. 3D, linear slopes [growth rate] in Fig. 3E) did not change with temperature (*PD*>0.15). However, temperature did affect the timing of initial spore production within hosts (i.e., intercepts of linear model). At the earliest point in the sacrifice series (day 8), spore load was highest at 26°C, intermediate at 32°C, and nearly zero at 20°C (Fig. 3D). Given thermally insensitive daily growth rates of parasites (*g*_*P*_; Fig. 3E), these head-starts were maintained over time (Fig. 3D). This effect on early spore production, coupled with host death rate (Fig 3C), explains the spore yield pattern. Final spore yield was lower at 32 than 26°C because there were fewer spores initially (on day 8) and hosts died more quickly (less time to produce spores). At 20°C, spore production started even later, but the delay was compensated for by much longer lifespans of infected hosts (lower death rate, *d*; Fig. 3C).

**Figure 3:**
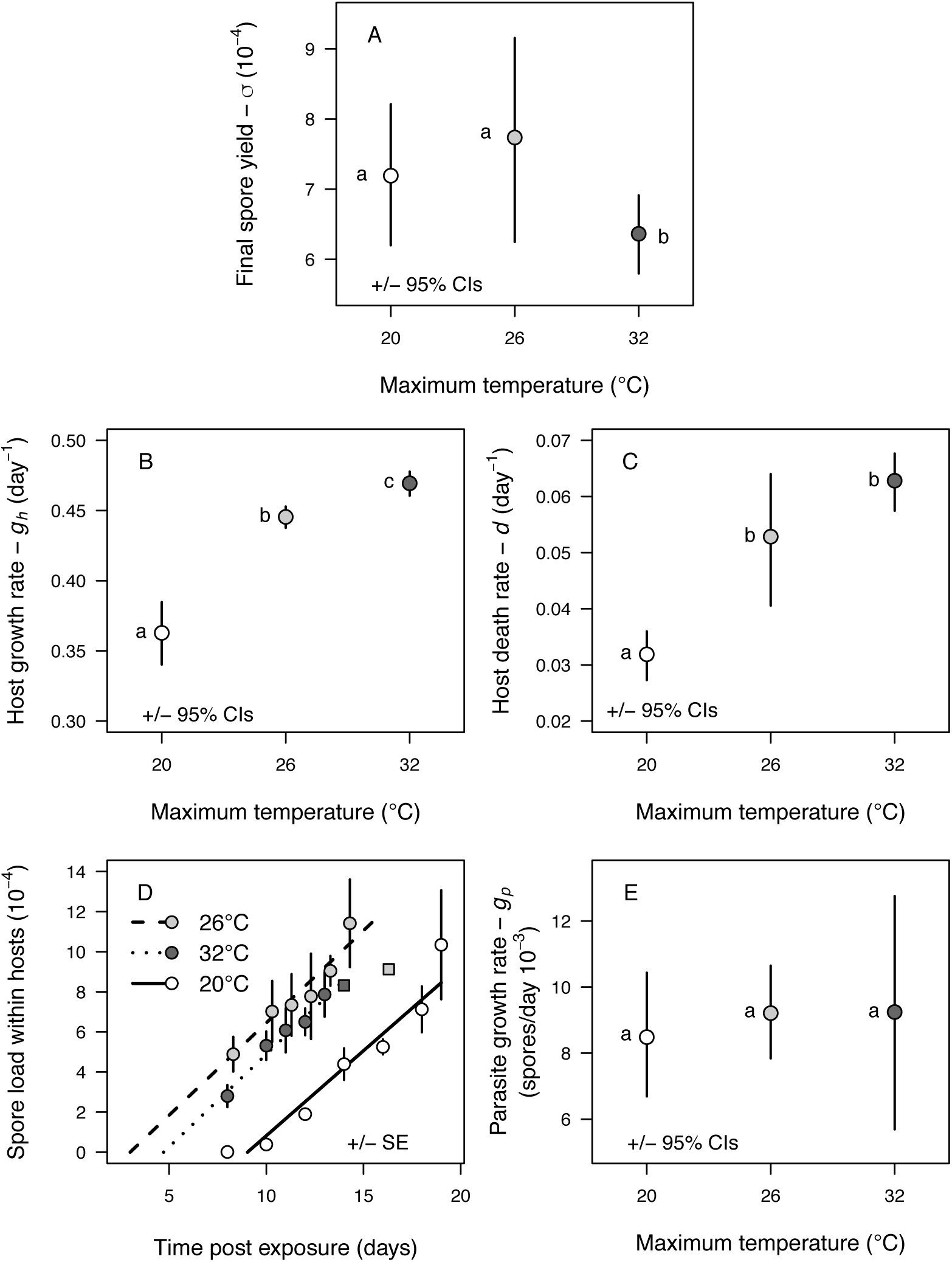
High temperature impacts on spore yield (*σ*, mechanism 3) and possible underlying traits. (A) Final spore yield at host death (*σ*) was lower at 32°C (dark grey) than at 20°C (white) or 26°C (light grey). (B) Host growth rate (*g*_*h*_) increased with temperature for all treatments. (C) Death rate of infected hosts (*d*) increased from 20 to 26°C and trended higher from 26 to 32°C. (D) Spore load within hosts through time at 32°C (dotted line), 26°C (dashed line), and 20°C (solid line), fit with linear models. 26°C points are shifted over for visual clarity of error bars. (E) Parasite growth rate (*g*_*p*_, slopes of lines in panel D) did not change with temperature. Hence, declining *σ* stems from higher death rate of infected hosts and low initial parasite growth, not slower growth rates of hosts or parasites. (A-C,E) Error bars show 95% CIs.(D) Error bars show SE; square points are single hosts. Letters indicate significant differences.

### Mechanisms 4 & 5: rearing (*ρ*) and free-living spore (*φ*) effects on infectivity

Spore infectivity (measured as transmission rate) responded unimodally to temperature in the previous infection (rearing effect on spore quality, *ρ*; Fig. 4A). Infectivity increased significantly for spores made at 20 versus 26°C for one of two spore sources (*p*=0.0083 for spores from *β* + *u* measurement assay [square, Fig 4A]; *p*=0.092 for spores from within-host growth assay [diamond]). Infectivity then declined for spores made at 26 versus 32°C (*p*=0.0001 for both spore sources). Infectivity was significantly lower for spores made at 32 versus 20°C for one of two spore sources (*p*=0.16 for spores from *β* + *u* measurement assay [square]; *p*=0.026 for spores from within-host growth assay [diamond]). The parameter *ρ* (Fig. 4C) shows the rearing effect pooled for both spore sources and normalized by transmission rate at 20°C (used for calculating transmission potential).

**Figure 4:**
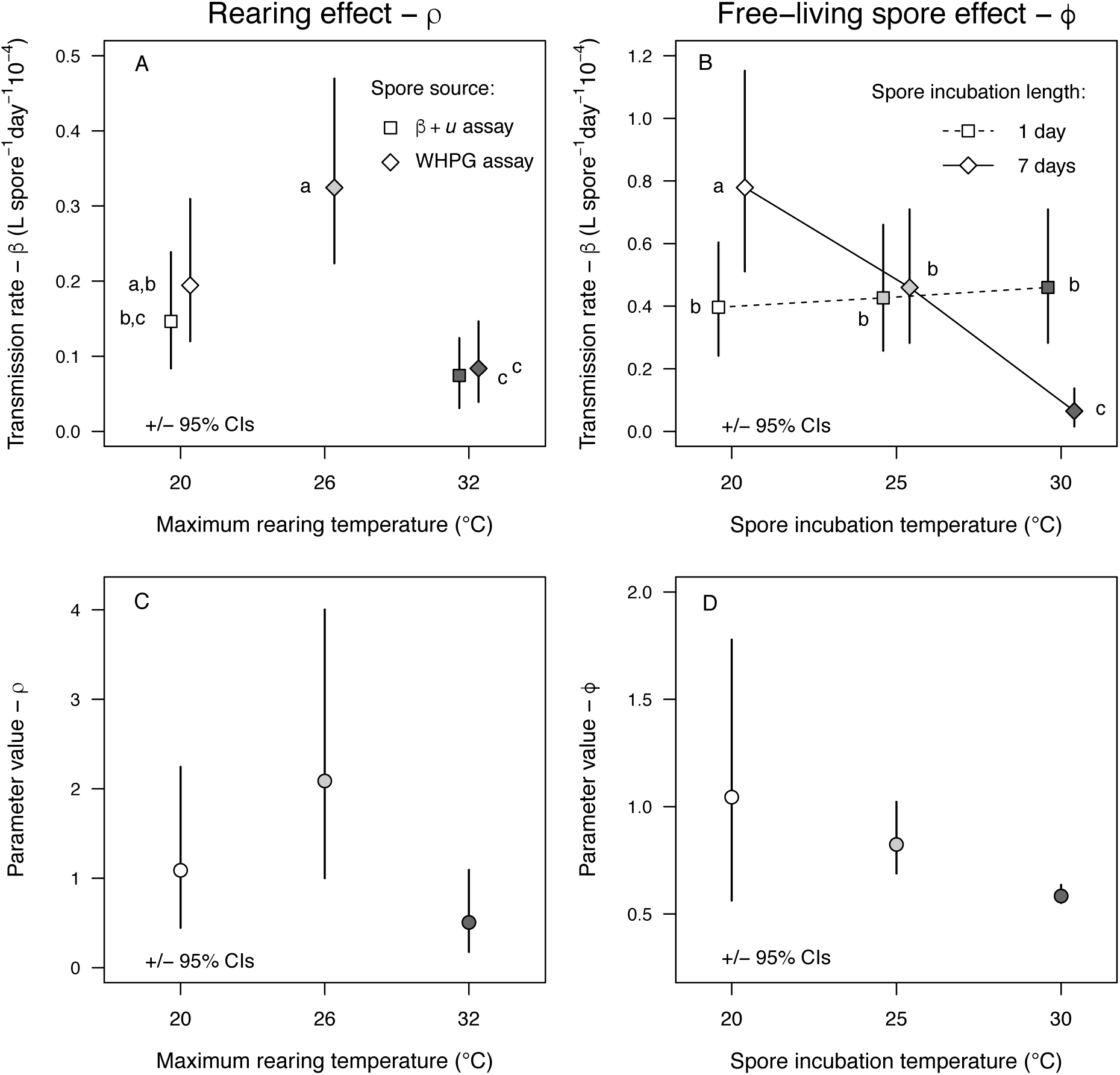
High temperature impacts on a rearing effect (ρ, mechanism 4) and harm to free-living spores (φ, mechanism 5). Variation in transmission rate from common garden infection assays reflects differences in spore infectivity. (A) Spores came from the β + *u* measurement assay (Fig. 2; squares) and the within host parasite growth assay (‘WHPG’; Fig. 3; diamonds). Spore infectivity increased with rearing temperature from 20°C (white) to 26°C (light grey; β + *u* only) and decreased with rearing temperatures from 26°C to 32°C (dark grey, both spore sources). Spore infectivity was lower at 32°C than at 20°C (WHPG spores only). (B) Spore infectivity decreased when free-living spores were incubated in high temperatures for 7 days but not for 1 day. Storage at 4°C for 6 days (for all 1-day incubations) also lowered spore infectivity relative to the 7-day incubation at 20°C. (C-D) Parameter values (transmission rate scaled by values at 20°C) for (C) rearing effect, ρ, and (D) free-living effect, *φ*. Phi values also based on time-weighted model (see text for details). Error bars show 95% CIs. Letters indicate significant differences.

The thermal environment of free-living spores also impacted their infectivity (*φ*; Fig 4B,D). Spore infectivity decreased with higher incubation temperatures after 7 days (20 versus 25°C: *p*=0.0031; 25 versus 30°C: *p*<0.0001; diamonds on Fig. 4B). However, spore infectivity did not change after 1-day incubations (flat line in Fig. 4B [squares]; 20 versus 25°C: *p*=0.65, 25 versus 30°C: *p*=0.64). All 1-day incubations used stored (refrigerated) spores. They had lower infectivity than the 7-day incubation at 20°C, likely because storage at 4°C also lowers spore infectivity (1 versus 7-day incubations at 20°C: *p*<0.0001) (Duffy & Hunsberger, 2019). The parameter *φ* (Fig. 4D) shows the free-living spore effect assuming that spores lose infectivity gradually over seven days as they are consumed by hosts (see *Methods* and Appendix) and normalized by transmission rate at 20°C (used for calculating transmission potential).

### Transmission potential (*f u σ ρ φ*)

Transmission potential, the product of all five mechanisms (*f u σ ρ φ*), responded unimodally to high temperatures. This metric first increased from 20 to 25/26°C (*PD*=0.017); then, it decreased from 25/26 to 30/32°C (*PD*=0.0001; Fig. 5A, ‘full transmission potential’). Transmission potential at 30/32°C trended (non-significantly) lower than at 20°C (*PD*=0.11). Thus, high temperatures do not constrain disease enough via these five mechanisms to explain the absence of summer epidemics.

**Figure 5:**
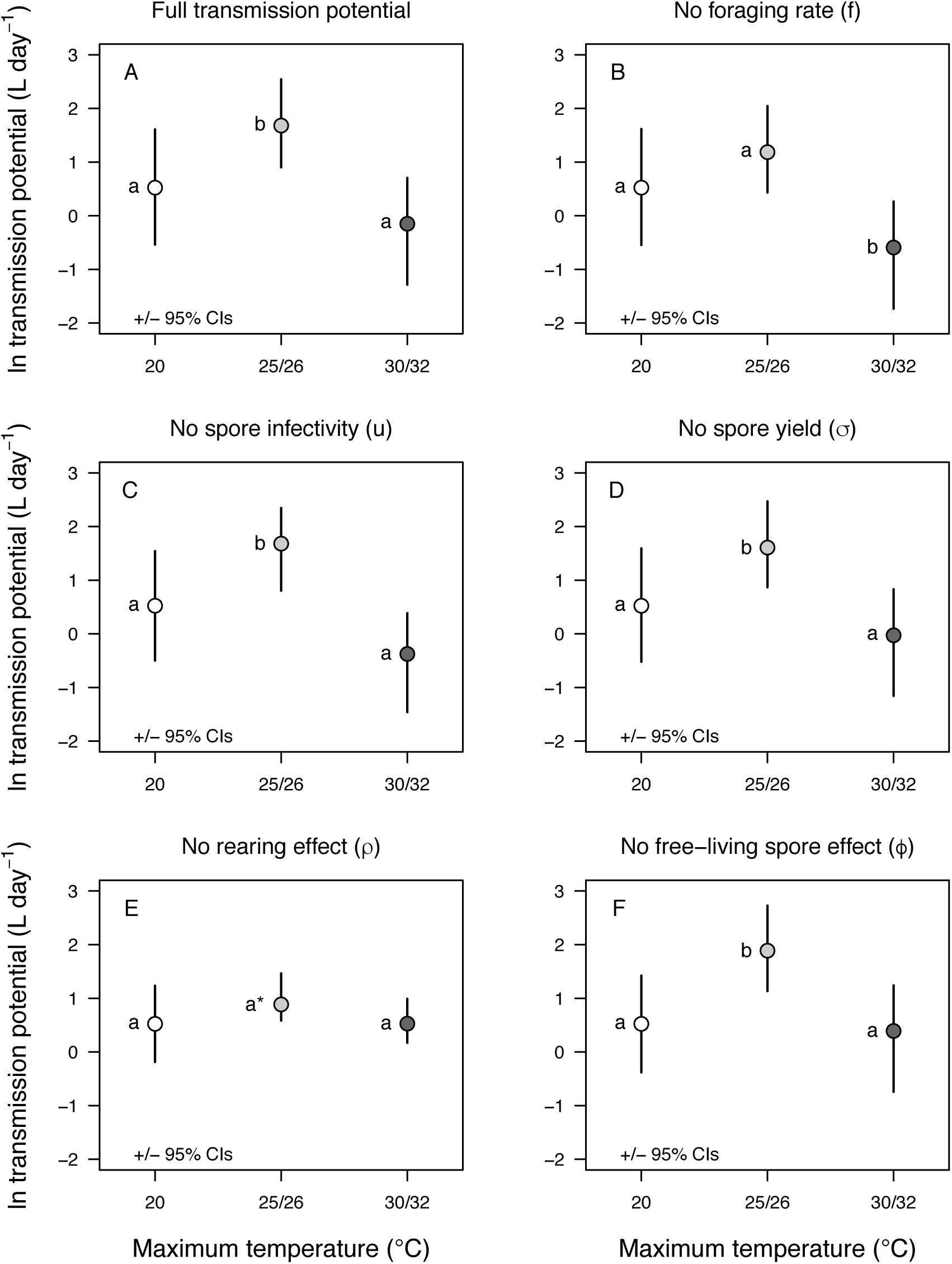
High temperature impacts on transmission potential. (A) Transmission potential (*fuσρφ*) responds unimodally, increasing from 20°C (white) to 25/26°C (light grey) and decreasing from 25/26°C to 30/32°C (dark grey). (B-F) Transmission potential with each mechanism held constant to show sensitivity to each parameter: (B) foraging rate (*f*), (C) spore infectivity from within-host effects (*u*), (D) spore yield (*σ*), (E) rearing effect (ρ), and (F) harm free-living spores (*φ*). The rearing effect (E) has the largest impact on transmission potential (hence, without it, the response of transmission potential is flat with temperature). Error bars show 95% CIs. Letters indicate significant differences. Y-axis is ln-transformed.

The initial increase in transmission potential from 20 to 25/26°C was driven most strongly by host foraging (*f*, mechanism 1) and the rearing effect on spore quality (*ρ*, mechanism 4): holding either trait constant removes the significant difference between temperatures (Fig. 5B and 5E, respectively). The subsequent drop in transmission potential from 25/26 to 30/32°C was driven most strongly by the rearing effect (*ρ*): holding it constant again removes the significant difference (Fig. 5E). Harm to free-living spores (*φ*, mechanism 5) also contributes somewhat (Fig. 5F vs. Fig. 5A), though not enough to affect the statistical significance. Additionally, the thermal response of host foraging (*f*) is key for maintaining transmission at high temperatures: without increased exposure to spores, the remaining mechanisms would significantly reduce transmission at 30/32°C compared to 20°C (Fig 5B). Spore infectivity from within-host processes (*u*, mechanism 2) and spore yield (*σ*, mechanism 3) had no effect (Fig. 5C vs. Fig. 5A) and very little effect (Fig. 5D vs. Fig. 5A) on transmission potential, respectively.

## DISCUSSION

We investigated upper thermal constraints on fungal epidemics in a *Daphnia* zooplankton host. The seasonality of the autumnal epidemics suggested that hot conditions might constrain disease: epidemics usually start after lakes cool from maximal summer temperatures in the epilimnion (29-32°C). We tested five potential thermal constraints on transmission. First, foraging (exposure) rate of hosts (*f*) increased at high temperatures (Fig 2B), while, second, high temperatures did not affect the infectivity of spores from within-host processes (*u*; Fig 2C). Thus, high temperatures increased transmission rate, *β* (where *β=uf*; Fig 2A). Third, spore yield (*σ*) declined slightly at 32°C (Fig 3A). Fourth, a rearing effect on spore quality driven by temperature during the previous infection (*ρ*) emerged: spores made at 32°C were less infectious than those made at 26°C and sometimes 20°C (for one of two spore sources, Fig 4A). Finally, harm to free-living spores (*φ*) lowered infectivity as temperature increased (Fig 4B). Overall, transmission potential is much lower at 32°C than 26°C, but still similar to at 20°C (Fig 5A), a temperature that easily supports epidemics in both nature (Shocket, Strauss, et al., 2018) and laboratory environments (Civitello et al., 2012; Shocket, Strauss, et al., 2018). Thus, maximally high temperatures do constrain disease, but not sufficiently to explain the absence of summer epidemics on their own.

Contrary to our initial hypothesis, high temperatures *increased* transmission rate (Fig 2A). In principle, high temperatures can lower infection success if pathogens tolerate heat less well than hosts (Thomas & Blanford, 2003). For instance, many ectothermic hosts behaviorally induce fever to reduce the negative costs of infection (Rakus et al., 2017; Stahlschmidt & Adamo, 2013). Further, fungi are particularly sensitive to high temperatures compared to other pathogen taxa (Robert & Casadevall, 2009) and fungal pathogens are often limited by high temperatures (Berger et al., 2004; Carruthers et al., 1992; Langwig et al., 2015; Raffel et al., 2010; Thomas & Blanford, 2003). However, high temperatures did not interfere with this fungus’s success at either stage of transmission: the day of exposure, when most spores penetrate the host’s gut, or infection establishment, when the fungus replicates and develops within the host (Stewart Merrill & Cáceres, 2018). Instead, high temperatures elevated host foraging rate (Fig 2B), which increases exposure to parasites, thereby increasing transmission rate (Hall et al., 2007). In lakes, the thermal response of foraging (exposure) drives variation in the size of epidemics, which occur in autumn: epidemics that start earlier in warmer conditions grow larger than those starting later and colder (Shocket, Strauss, et al., 2018). This foraging-controlled exposure to parasites is a potentially general mechanism: higher temperatures also increase outbreak size for armyworms that consume virus particles on leaves (Elderd & Reilly, 2014). However, transmission plateaued with temperature for another ingested *Daphnia* pathogen (Vale, Stjernman, & Little, 2008).

Spore yield (*σ*) declined at the highest temperature (32°C; Fig. 3). Although the effect on transmission potential was minimal (Fig 5D), the results for related traits provide mechanistic insights into host-parasite interactions. Parasite burdens often decline at temperatures near the thermal maxima of the host and/or parasite, e.g., for nematodes in slugs (Wilson, Digweed, Brown, Ivanonva, & Hapca, 2015), trematodes in snails (Paull, Lafonte, & Johnson, 2012), bacteria in *Daphnia* (Vale et al., 2008) and fruit flies (Lazzaro, Flores, Lorigan, & Yourth, 2008), and powdery mildew in plants (Laine, 2007). In theory, reduced parasite production at hot temperatures could arise from several mechanisms. First, parasite production could decline if host growth slows, since spore yield often scales with host growth, at least along resource gradients (Hall, Knight, et al., 2009; Hall, Simonis, et al., 2009). However, here host growth rate (*g*_*h*_) increased with temperature while spore yield was flat and then decreased (Fig 3B).

Therefore, spore production was decoupled from host growth rate (i.e., the link between host growth and parasite production that occurs for resources did not occur for temperature). Second, the parasite itself could grow more slowly at high temperatures. For example, high temperatures slow bacterial growth inside fruit flies (Lazzaro et al., 2008), fungal growth in grasshoppers (Springate & Thomas, 2005), and fungal growth on warm-adapted (but not cold-adapted) amphibians (Cohen et al., 2017). In contrast, here parasite growth rate (*g*_*p*_) did not respond to temperature (slope in Fig 3D; Fig 3E).

Instead, the decline in spore production at high temperatures arose from a combination of host death rate and the timing of initial spore production. Temperature determined spore load on day 8 (the earliest sampling time in the assay; Fig. 3D). Based on that information (and the constant parasite growth rates, Fig 3E), we infer that spore production began earliest at 26°C, followed by 32°C, and then 20°C. These head starts were maintained over time and explain the spore yield pattern when combined with death rate of infected hosts (Fig. 3C). In general, shorter lifespan of infected hosts decreases time for spore production, thereby depressing spore yield (Auld et al., 2014; Civitello et al., 2012). Here, spore yield was lower at 32 than 26°C because spore production started later and hosts died more quickly. At 20°C, spore production started even later, but longer host lifespan compensated for this delay (i.e., the fungus had longer to grow within hosts). Do similar patterns exist in other systems? Unfortunately, few studies focus on traits underlying thermal responses of parasite load. Hence questions remain: How often does temperature change the timing versus the rate of parasite production? How often does temperature decouple positive relationships between host growth and parasite production? The answers matter because spore yield can influence epidemic size for obligate killer parasites (like the fungus here: Civitello et al., 2015). Thus, developing a general framework from data across host-parasite systems remains a fruitful area for future research.

High temperatures reduced transmission potential via two effects on spore infectivity that act outside the focal host. First, a rearing effect on spore quality (*ρ*) driven by temperature of spore production in the previous host elevated (26°C) and then lowered (32°C) spore infectivity (compared to 20°C). Rearing effects on parasite performance can arise with variation in resources consumed by hosts (Cornet, Bichet, Larcombe, Faivre, & Sorci, 2014; Little, Birch, Vale, & Tseng, 2007; Tseng, 2006), temperature experienced by hosts (Shocket, Vergara, et al., 2018), or host genotype (Searle et al., 2015). These understudied rearing effects may drive performance of parasites to an unappreciated extent (Shocket, Vergara, et al., 2018). Second, harm to free-living spores (*φ*, including spore mortality) also inhibited infection at high temperatures. After seven days in 30°C, spores lost 92% of their initial infectivity. This constraint may arise in other systems: for example, high temperatures elevate mortality in free-living helminths of Arctic ungulates (Molnár et al., 2013). However, in the planktonic system here, the 7-day result likely exaggerates the thermal constraint. While difficult to quantify, physical sinking, consumption (Civitello, Pearsall, Duffy, & Hall, 2013; Penczykowski, Hall, et al., 2014; Shocket, Vergara, et al., 2018; Strauss, Civitello, Cáceres, & Hall, 2015) and damage from radiation (Overholt et al., 2012) likely remove most spores before 7 days. To acknowledge this mortality, we weighted this component of infectivity (*φ*) using a model of spore longevity. Assuming this modeled weighting reflects reality in lakes, the free-living effect lacks enough strength to inhibit epidemics during summer, even when combined with the other mechanisms (see Appendix for sensitivity analysis of the time-weighting model). However, more realistic dynamical models and better resolved trait data for the free-living spore effect could change the estimates for how high temperatures affect transmission.

Although the impact of temperature on these five mechanisms does not explain the lack of epidemics during summer, other co-varying environmental factors could combine with thermal effects to sufficiently inhibit transmission. Such factors include damage to free-living spores by solar radiation (Overholt et al., 2012), consumption of spores by resistant zooplankton species that are more abundant earlier in the year (Penczykowski, Hall, et al., 2014), and low spore production due to poor quality of host food resources (Hall, Knight, et al., 2009). These mechanisms could contribute to the observed field pattern, and interact with the thermal effects examined here. Furthermore, climate change could disrupt covariation among drivers. For example, high temperatures may persist later in the year when damaging solar radiation is less intense. Incorporating these other factors may help explain the current field pattern and improve predictions for how climate change will impact epidemics. These predictions should also explicitly account for the effects of temperature variation and extremes, which have distinct impacts on organismal performance (Dowd, King, & Denny, 2015). Here, we employed a relevant form of thermal variation, mimicking migratory behavior of hosts in stratified lakes, but did not isolate effects of thermal variation. Future efforts could estimate this effect to better predict how climate change will impact the host, the parasite, and their interaction.

The current prevailing view argues that hot temperatures should constrain disease transmission in nature (Altizer et al., 2013; Lafferty, 2009; Lafferty & Mordecai, 2016). This constraint arises when unimodal thermal reaction norms depress key traits that drive disease spread. However, such constraints have been rigorously tested in only a handful of systems. Here, we hypothesized that high summer temperatures limit transmission of a zooplankton-fungus disease system with autumnal epidemics (i.e., during cooler conditions). High temperatures constrained disease transmission enough to produce a unimodal thermal response. This response arose primarily through a rearing effect on spore quality and due to harm to free-living spores. However, the thermal mechanisms estimated here were not sufficient explain the lack of summer epidemics. Hence, we draw two major lessons. First, we need to continue to rigorously evaluate multiple mechanisms of thermal constraints on components of disease transmission. Second, our example cautions against drawing conclusions about constraints on disease from warming based on field patterns and theory alone.

## ACKNOWLEDGMENTS

K. Boatman (2009 and 2010), Z. Brown and K. Malins (2011), O. Schmidt (2013), A. Bowling (2014), and P. Orlando and J. Walsman (2015) conducted field sampling. MSS was supported by NSF GRFP. This work was supported by NSF DEB 0841679, 0841817, 1120316, 1120804, 1353749, and 1354407.

## SUPPORTING INFORMATION

Additional supporting information may be found in the online version of this article.

Appendix S1: Methods, Figures, and Tables

Figure S1: Components of a simple model used to estimate parameter *φ*.

Figure S2: Sensitivity analysis for spore consumption model parameter (*c*) affecting damage to free-living spores (*φ*) and transmission potential.

Table S1: Raw results from the *β + u* measurement assay.

Table S2: Sample sizes for estimating spore yield (*σ*) and related traits.

Table S3: Raw results from the rearing effect (*ρ*) and free-living spore effect (*φ*) assays.

Table S4: *p*-values from randomization tests.

Table S5: *p*-values and ΔAIC from model selection.

Table S6: PD (probability density) values for traits.

Table S7: PD (probability density) values for transmission potential ‘sensitivity analysis’ calculations.

