## Supplemental Methods, Figures, and Tables for "Can hot temperatures limit disease transmission? A test of mechanisms in a zooplankton-fungus system"

**APPENDIX S1: Methods, Figures, and Tables**

**List of Contents**

1. Experimental Methods and Parameter Calculations
   a. General Methods
   b. Mechanism 1: Foraging rate (*f*)
   c. Mechanism 2: Transmission rate (*β*) and per spore infectivity (*u*)
   d. Mechanism 3: Spore yield (*σ*) and other measures of host and parasite growth – growth rate of hosts (*g_h_*), growth rate of parasites within hosts (*g_p_*), death rate of infected hosts (*d*)
   e. Mechanisms 4 & 5: rearing (*ρ*) and free-living spore (*φ*) effects on infectivity
   f. Transmission Potential Calculations
2. Statistical Methods and Results of Pairwise Comparison Tests
   a. Calculating PD values from bootstrapped parameter distributions
   b. Table S4: *p*-values from randomization tests (*β*, *u*, *ρ*, *φ*, and *d*)
   c. Table S5: *p*-values and ΔAIC from model selection (*σ*) and t-tests (*g_h_*)
   d. Table S6: PD-values for main parameters (*f*, *g_p_*, and transmission potential)
   e. Table S7: PD-values for ‘sensitivity analysis’ of transmission potential
3. References

**Experimental Methods and Parameter Calculations**

**General Methods**

All experiments used a single clonal genotype of the *Daphnia dentifera* zooplankton host isolated from a lake in Michigan (‘Standard’). Hosts were kept in filtered lake water and fed 1.0 mg dry mass / L nutritious green algae (*Ankistrodesmus sp.*) daily.

Mechanism 1: Foraging rate (*f*)

*Foraging rate* (*f*): The experiment produced data for 37 hosts ranging from 0.92 to 1.75 mm in size (average = 1.39 mm). Hosts were acclimated to fluctuating exposure to 30°C (8 hours at 20°C and 16 hours at 30°C) for one week prior to the experiment. Individual hosts were transferred into centrifuge tubes containing 15 ml of water and 1.0 mg dry mass / L algae. The tubes were then placed at 30°C, kept dark, and inverted every 30 minutes to resuspend algae. After grazing for 8.5 hours, hosts were measured (for body length: eye to base of tail spine at 50X). We measured *in vivo* fluorescence of the remaining algae and compared it to ungrazed controls (using a Turner Trilogy Laboratory Fluorometer).

We assumed there was no algal growth during the dark assay. Thus, with only a single susceptible host (*S*) per tube, the differential equation for algae (modified from eq. 3e) becomes:

$\frac{dA}{dt}= -fSA$	eq. S1

where *f* is the foraging rate in this linear functional response. Solving this exponential equation for resource density (*A*) yields:

$A_{rem}= A_{0}e^{-fSt}$	eq. S2

where *A_r_*_em_ is the remaining algae, *A*_0_ is the initial amount of algae (at time *t* = 0), *S* is the density of hosts (i.e., 1/*V*, where *V* is the experiment volume), and *t* is the duration of the trial (~8.5 hours). For each individual host, we calculated *f* using log-transformed version of eq. S2:

$f=\frac{ln(A_{0}/A_{rem})}{St}$	eq. S3

With these estimates of *f* and body length measurements for each host*,* we used maximum likelihood estimation to fit a normally-distributed function for *f* at 30°C according to:

$f=\hat{f}L^{\gamma}$	eq. S4

Where $\hat{f}$ is the size-specific foraging rate, *L* is body length (in mm), and *γ* is the power coefficient for size. The estimated value for *γ* was 2.56 (95% CIs: 1.85–3.71).

Mechanism 2: Transmission rate (*β*) and per spore infectivity (*u*)

*Transmission* (*β*): In the *β* + *u* measurement assay, for each of the four treatments, 49 six-day-old hosts (all acclimated at 32°C for two generations) were exposed individually (in 15 ml tubes) to a fixed dose of spores (45 sp/ml) at 20°C or 32°C (‘exposure temperature’). The next day, hosts were moved to individual beakers containing 80 ml of fresh water, and one group from each exposure temperature was moved to the other thermal environment (‘infection establishment temperature’). Hosts were checked daily to see if they were alive and moved to fresh water every other day. All dead hosts were collected beginning on day 7. On day 11 for the 32°C infection establishment groups or day 13 for the 20°C infection establishment groups we visually diagnosed all living hosts with a dissecting scope (20-50X magnification), and discarded the uninfected hosts. We determined infection prevalence of the previously deceased hosts during spore quantitation (see *Mechanism 3*). The raw results are given below in Table S1.

We estimated transmission rate (*β*) for this experiment and others described below using maximum likelihood (with the ‘bbmle’ package in R). We based our estimation on a simplified transmission model (Bertram, Pinkowski, Hall, Duffy, & Cáceres, 2013; Hall, Tessier, Duffy, Huebner, & Cáceres, 2006), where susceptible hosts (*S*) are lost (and become infected) after contacting spores (Z) with transmission rate *β*:

$\frac{dS}{dt}=-\beta ZS$	eq. S4

We solve this equation for the remaining susceptible hosts (*S_f_*) after exposure time *t_e_* yielding:

$S_{f}=S_{0} e^{-\beta Zt_{e}}$	eq. S5

where *S*_0_ is the initial numbers of hosts in a beaker (here, always one). The probability of remaining uninfected (*p_u_*) equals:

$p_{u}= e^{-\beta Zt_{e}}$	eq. S6

We then use a binomial distribution to model the number of uninfected hosts in each tube.

In the two 32°C exposure treatments (32/32°C and 32/20°C), infection prevalence was very high (only 6/47 and 3/45 animals remained uninfected, respectively). Therefore, some of the bootstrapped samples resulted in 100% infection prevalence. In these cases, the estimate of transmission rate (*β* = 1.5·10^-3^) is unrealistically high by an order of magnitude. Therefore, we corrected these estimates of *β* using the likelihood profile method (Cole, Chu, & Greenland, 2014). This approach calculates the likelihood profile to find the parameter (here, *β*) value where the loglikelihood is 1.92 units lower than the maximum value, i.e., the upper limit of the confidence interval (1.92 is half the 3.84 chi-square value for 1 degree of freedom). We calculated this upper limit for the higher prevalence treatment (32/20°C) and substituted the resulting value (*β* = 9.05·10^-5^) for the bootstrapped samples with 100% prevalence in both treatments.

**Table S1: Raw results from the *β + u* measurement assay.** Total hosts reflects the number surviving past day 7 when it becomes possible to diagnose infection status.

| **Treatment Group**  **(Exposure / Infection Establishment)** | **Infected hosts / Total Hosts** |
| --- | --- |
| 20/20°C | 30 / 46 |
| 32/20°C | 42 / 45 |
| 20/32°C | 24 / 49 |
| 32/32°C | 41 / 47 |

Mechanism 3: Spore yield (*σ*) and other measures of host and parasite growth

*Spore yield* (σ): We measured spore yield from hosts from two assays: the *β + u* measurement assay (with only 20/20 and 32/32°C exposure / infection establishment temperature treatments; see above, *Mechanisms 1 & 2*) and the within host parasite growth assay (20, 26, and 32°C; see below). Sample sizes are given below in Table S2. Spore yield did not differ between the same temperature groups in the different assays (*p* = 0.65 for 20°C, *p* = 0.93 for 32°C), so we pooled data from like treatments. Hosts were homogenized to estimate spore yield for each individual (counted at 200X with a hemocytometer).

*Growth rate of hosts* (g_h_): Twenty-four neonates were harvested from each temperature (20, 26, 32°C), housed and fed individually in 15 ml tubes with fresh water changed daily, then dried four days later. We calculated growth rate for each individual *i* (*g_hi_*):

$g_{h_{i}}=\frac{\ln\left( m_{i} \right)-\ln\left( \bar{m}_{0} \right)}{t}$	eq. S7

where *m_i_* is the mass of the individual, *m̄*_0_ is the average mass of the neonates (day 0) for that temperature (*N* = 8), and *t* is four days.

*Growth rate of parasites* (g_p_): Three groups of 48–51 six-day-old hosts (all acclimated at 32°C for two generations) were exposed individually (in 15 ml tubes) to a fixed dose of spores (200 sp/ml) at room temperature (~20°C). The next day, we placed hosts into individual beakers (80 ml of water) and moved them to their thermal treatment (20°C, 26°C, and 32°C). We checked hosts daily for survival. Beginning on day 8 post-exposure, we sacrificed 7-10 hosts in each treatment every other day. After day 11 for 26°C and 32°C or day 14 for 20°C, we allowed the rest to die naturally. We counted spore load (in sacrificed hosts) or spore yield (in dead infected hosts).

With these spore load and spore yield data, we used linear models to estimate growth rate of parasite (*g_p_*) via a linear model for spores within hosts through time. We chose a linear response based the observed response in the data. Microbial growth is often modeled exponentially (log-linearly). Although we cannot do model comparison directly (i.e., with AIC scores) because the two sets of models are fit to different y-values, 1) the R^2^ values are higher for linear models (20°C: 0.73, 26°C: 0.12, 32°C: 0.29) than for the corresponding exponential models (20°C: 0.57, 26°C: 0.07, 32°C: 0.15) and 2) the model residuals are more normal (not shown). Additionally, the fungus here reproduces in a way that not typical for microbial populations, i.e., spores do not replicate to produce new spores. Rather, conidial cells replicate to produce new conidial cells, and then at some point conidial cells stop replicating and begin maturing into spores instead.

We fit separate models for each temperature treatment. The models for 20°C and 32°C yielded biologically sensical interpretations. However, the model for 26°C did not: the x-intercept (i.e., mean day of first spore production) was close to 1.5 days, which is unrealistically early for spore production. Thus, we fixed the x-intercept to a more reasonable value (3 days). This model fit the data similarly well (ΔAIC = -1.8, with the fixed-intercept model having the lower AIC score since it had one fewer parameter; thus, the parameter-uncorrected difference in fits was only -0.2 AIC units).

*Death rate of parasites* (d): We assumed that time until death followed an exponential distribution (McCallum, 2000) and estimated the death rate using maximum likelihood (package ‘bbmle’ in R). The exponential distribution provides a likelihood function for the constant death rate, *d,* given the time-until-death data for each focal uninfected host, *t_d_*:

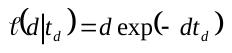
.	eq. S8

(All animal died in this dataset, obviating the need to include censored individuals).

**Table S2: Sample sizes for estimating spore yield (***σ***) and related traits.** *β + u* measurement assay numbers only include hosts from the 20/20°C and 32/32°C treatments who became infected. WHPG = within host parasite growth assay. Total hosts from WHPG assay (used to calculate growth rate of the parasite within hosts, *g_p_*) include parasite-killed hosts and sacrificed hosts; only parasite-killed hosts were included in the spore yield (*σ*) and death rate (*d*) calculations.

| **Temp. Group** | **Infected hosts *β + u* assay** | **Parasite-killed hosts WHPG assay** | **Total hosts WHPG assay** | **Total hosts for spore yield (***σ***) & death rate (d) calculations** |
| --- | --- | --- | --- | --- |
| 20°C | n = 30 | n = 14 | n = 47 | n = 44 |
| 26°C | NA | n = 26 | n = 45 | n = 26 |
| 32°C | n = 42 | n = 33 | n = 51 | n = 75 |

Mechanisms 4 & 5: rearing (*ρ*) and free-living spore (*φ*) effects on infectivity

To measure the rearing effect (*ρ*), for each spore treatment, we collected and pooled spores from 3-5 hosts that died from their infection within the same 3-day period. For the free-living spore effect (*φ*), we took a single batch of freshly homogenized spores and put 100,000 spores into six replicate dark amber plastic bottles containing 150 ml of filtered lake water. The bottles were covered loosely with aluminum foil to keep out light but allow airflow. Three bottles were immediately moved into the constant temperature environments. The other three bottles were kept at 4°C for six days, and then moved into the constant temperature environments for the final incubation day. (Standard procedure for storing spores is to collect infected animals as they die or are near death, and store them in Eppendorf tubes in a small volume of lake water at 4°C.) Then, to estimate both parameters, we used the spores (above) in two common garden infection assays at constant 20°C.

For each assay, either 45 (*ρ*) or 40 (*φ*) six-day-old hosts were exposed individually (in 15 ml tubes) to a fixed dose of spores (30 sp/ml) from each spore treatment. The next day, all hosts in a given treatment were moved to a single flask with fresh water. Ten days later, we diagnosed the infection status of the hosts and calculated the proportion of hosts infected for each treatment (and then calculated transmission rate using eq. S6, as described above). Initially uninfected animals were re-examined on day 14. The raw results are given below in Table S3.

When estimating the *φ* parameter, we used a simple model to account for the fact that spores are consumed over time. We assumed that spore infectivity declined linearly over time from day 1 to day 7 (Fig. A1B). (In the assay, there was no difference in infectivity after 1 day, and substantial differences after 7 days.) We assumed that spores are consumed by hosts at a constant rate, resulting an exponential decline in spores remaining in the environment over time (Fig. A1B). Then we multiplied the infectivity of spores on a given day by the proportion of spores consumed on that day. We parameterized the model (*y* = e^-0.25^*^x^*; where *y* = spores remaining and *x* = days) such that most spores (83%) were consumed by day 7, but a substantial number of spores still remain on day 7 itself (17%) and thus are weighted by the day 7 estimates of infectiousness. This choice moderates the effect on free-living spores to match our current understanding of the system without completely negating it. As a result, our measure of *φ* is more conservative (a smaller effect of high temperatures) than the transmission rate values from the 7-day incubations. It is very difficult to measure the persistence of spores in natural environments; however, to the best of our knowledge, most spores are consumed relatively quickly (Shocket et al., 2018). This simple model reflects our best guess for a reasonable set of spore dynamics. The development of better methods is needed to provide a more accurate estimate, but is beyond the scope of this paper.

We performed a sensitivity analysis to examine how our choice of parameter for the spore consumption model affects the results for transmission potential. We bound our possibilities for how to incorporate the free-living spore effect by having no effect (unreasonably weak influence) on one end and the full 7-day effect instantaneously applied to all spores (unreasonably strong influence) on the other. All parameter choices for the spore consumption model fall between these two options. The choice of parameter does have a small impact on the results, particularly for the 30/32°C treatment (Fig A2). However, the differences between the various parameter values are relatively small compared to the difference between using the spore consumption model with any parameter value and simply using the value for transmission rate after 7-day incubations (Fig A2C).

**Table S3: Raw results from the rearing effect (*ρ*) and free-living spore effect (*φ*) assays.** Hosts were infected using spores from the *β + u* measurement assay and within host parasite growth (WHPG) assay. Total hosts reflects the number surviving to day 10 when we diagnosed infection status.

| **Treatment Group** | **Infected hosts / Total Hosts** |
| --- | --- |
| *Rearing effect assay* |  |
| 20/20°C *β + u* spores | 16 / 45 |
| 32/32°C *β + u* spores | 9 / 45 |
| 20°C WHPG spores | 19 / 43 |
| 26°C WHPG spores | 28 / 45 |
| 32°C WHPG spores | 10 / 45 |
| *Free-living spore effect assay* |  |
| 20°C 1-day | 19 / 40 |
| 25°C 1-day | 19 / 38 |
| 30°C 1-day | 20 / 38 |
| 20°C 7-day | 28 / 39 |
| 25°C 7-day | 20 / 38 |
| 30°C 7-day | 4 / 40 |

**Figure S1: Components of a simple model used to estimate parameter *φ*.** (A) Over a 7-day period, transmission rate (reflecting spore infectivity) remains steady at 20°C but declines over time for 25 and 30°C. The magnitude of the decline increases with temperature. (B) Hosts consume spores at a constant rate, resulting in a negative exponential decline in the number of spores over time (*y* = e^-0.25^*^x^*; where *y* = spores remaining and *x* = days). The transmission rate (based on spore infectivity) on a given day is multiplied by the proportion of spores consumed by hosts on that day. Spores consumed after day 7 are assumed to have the 7-day value for transmission rate.

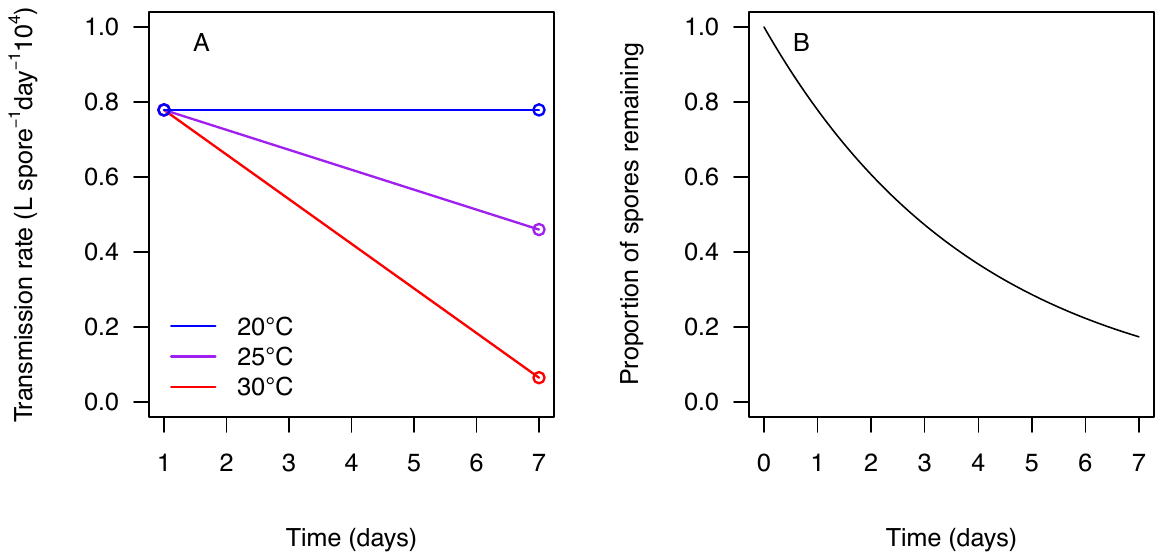

**Figure S2: Sensitivity analysis for spore consumption model parameter (*c*) affecting damage to free-living spores (*φ*) and transmission potential.** (A) The spore consumption model for four parameterizations: *c* = 0.5 (blue), 0.25 (cyan), 0.15 (orange), and 0.1 (red). (B) The resulting *φ* calculations, including the additional set where all spores are consumed on day 7 (black) with the appropriate estimated transmission rate (*β*). (C) The resulting transmission potential calculations, including the additional set where there is no damage to free-living spores (*φ* = 1, dark grey).

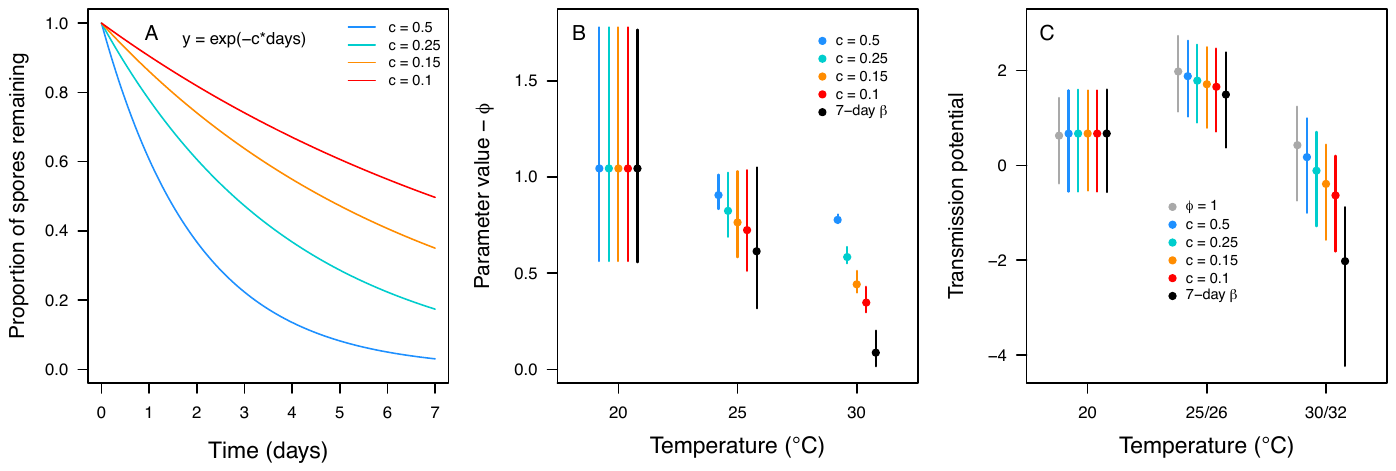

**Transmission Potential Calculations**

Spore infectivity (*u*) came from the 20/20 and 32/32°C treatments of the *β + u* measurement assay. Since this experiment lacked a 25 or 26°C treatment, we took the mean infectivity of the 20/20°C and 32/32°C treatments. Although they differed slightly, the values for *u* were not significantly different (*p* = 0.37). Because we lacked an intermediate temperature treatment (high temperature = 26°C) for the *β + u* measurement assay, it is possible that *u* responds unimodally to temperature, but we were unable to detect that response. Thus, we may be underestimating the transmission potential for the intermediate temperature treatment. We chose not to pursue this experiment due to limited time and resources, as it was not critical for our main question.

**Statistical Methods and Results of Pairwise Comparison Tests**

For *f* and transmission potential, we used the bootstrapped distributions to compare treatments. Specifically, we calculated the cumulative probability density of the best estimate from one treatment according to the distribution of the other. Thus, each comparison yields two probability-density values (because the comparison goes both ways: which treatment provides the best estimate and which provides the distribution). These PD-values are analogous to *p*-values, but bounded: 0<PD<0.5 (instead of 0<*p*<1). If *PD*=0, the best estimate is more extreme than all values in the bootstrapped distribution; if *PD*=0.5, the best estimate falls in the center of the distribution. We considered treatments significantly different if *PD*<0.025 (equivalent to a two-tailed test). Although the two values for a given treatment-comparison differed quantitively, they always agreed with each other regarding the significance threshold, with one exception. This exception was the sensitivity analysis where we calculated transmission potential without including the rearing effect (*ρ*). In that case, the best estimate for 25/26°C was within the 95% CIs for the other temperature treatments, but the best estimates for 20 and 30/32°C were not within the 95% CI for 25/26°C.

**Table S4: *p*-values from randomization tests.** Parameters tested are transmission rate (*β*), spore infectivity (*u*), death rate of infected hosts (*d*), the rearing effect (*ρ*), and harm to free-living spores (*φ*). Tests used 10,000 samples.

| **Parameter** | **Treatment Contrast / Model** | **p-value** |
| --- | --- | --- |
| Transmission rate (*β*)*^1^* | 20°C Exp. vs. 32°C Exp. (20°C Dev.) | ***p* = 0.0007** |
| (Fig 2A) | 20°C Exp. vs. 32°C Exp. (32°C Dev.) | ***p* = 0.0001** |
|  | 20°C Dev. vs. 32°C Dev. (20°C Exp.) | *p* = 0.10 |
|  | 20°C Dev. vs. 32°C Dev. (32°C Exp.) | *p* = 0.31 |
|  | 20°C Exp. & Dev. vs. 32°C Exp. & Dev. | ***p* = 0.0068** |
| Spore infectivity (*u*)*^1^* | 20°C Exp. vs. 32°C Exp. (20°C Dev.) | ***p* = 0.034** |
| (Fig 2B) | 20°C Exp. vs. 32°C Exp. (32°C Dev.) | ***p* = 0.0052** |
|  | 20°C Dev. vs. 32°C Dev. (20°C Exp.) | *p* = 0.10 |
|  | 20°C Dev. vs. 32°C Dev. (32°C Exp.) | *p* = 0.31 |
|  | 20°C Exp. & Dev. vs. 32°C Exp. & Dev. | *p* = 0.37 |
| Death rate of infected hosts | 20°C vs. 26°C | ***p* < 0.0001** |
| (Fig 3B) | 26°C vs. 32°C | *p* = 0.063 |
|  | 20°C vs. 32°C | ***p* < 0.0001** |
| Rearing effect (*ρ*)*^2^* | WHPG 20°C spores vs. 26°C | *p* = 0.092 |
| (Fig. 4A) | *β* + *u* 20°C spores vs. 26°C | ***p* = 0.0083** |
|  | 26°C vs. WHPG 32°C spores | ***p* = 0.0001** |
|  | 26°C vs. *β* + *u* 32°C spores | ***p* = 0.0001** |
|  | 20°C vs. 32°C, both WHPG spores | ***p* = 0.026** |
|  | 20°C vs. 32°C, both *β* + *u* spores | *p* = 0.16 |
| Free-living spore effect (*φ*)*^3^* | 20°C vs. 25°C (1 day) | *p* = 0.65 |
| (Fig. 4B) | 25°C vs. 30°C (1 day) | *p* = 0.64 |
|  | 20°C vs. 30°C (1 day) | *p* = 0.50 |
|  | 20°C vs. 25°C (7 day) | ***p* = 0.0031** |
|  | 25°C vs. 30°C (7 day) | ***p* < 0.0001** |
|  | 20°C vs. 30°C (7 day) | ***p* < 0.0001** |
|  | 20°C (1 day vs. 7 day) | ***p* < 0.0001** |

*^1^* The *β* + *u* measurement experiment used factorially crossed exposure (Exp) and infection development (Dev) treatments. *^2^* Rearing effect (*ρ*) was estimated using spores from two experiments, the *β* + *u* measurement assay (‘*β* + *u*’; Fig. 2) and the within-host parasite growth assay (‘WHPG’; Fig. 3). *^3^* Comparisons are only within 1 day temperature treatments and 7 day treatments, except for a planned contrast between 1 and 7 day at 20°C.

**Table S5: *p*-values and ΔAIC from model selection**. ΔAIC from model selection for (spore yield, *σ*) and *p*-values from t-tests (host growth rate, *g_h_*).

| **Parameter** | | **Treatment Contrast / Model** | **p-value/ΔAIC (Aikaike Weight)** |
| --- | --- | --- | --- |
| Model competition | |  |  |
| Spore yield (*σ*)*^1^*  (Fig. 3A) | 2 means (32°C vs. 20 & 26°C), 2 sd (32°C vs. 20 & 26°C) | | *Winning model^2^* (0.40) |
|  | 2 means (32°C vs. 20 & 26°C), 3 sd | | ΔAIC = 1.53, *p* = 0.49 (0.19) |
|  | 3 means, 2 sd (32°C vs. 20 & 26°C) | | ΔAIC = 1.62, *p* = 0.54 (0.18) |
|  | 1 mean, 2 sd (32°C vs. 20 & 26°C) | | ΔAIC = 2.00, ***p* = 0.045 (0.15)** |
|  | 3 means, 3 sd (full model) | | ΔAIC = 3.17, *p* = 0.66 (0.08) |
|  | 2 means (32°C vs. 20 & 26°C), 1 sd | | ΔAIC = 7.63, ***p* = 0.0019 (0.009)** |
|  | 1 mean, 1 sd (null model) | | ΔAIC = 9.73, ***p* = 0.0010 (0.003)** |
| T-tests |  | |  |
| Growth rate of hosts (*g_h_*) | 20°C vs. 26°C | | ***p* = 4.7 x 10^-6^** |
| (Fig. 3B) | 26°C vs. 32°C | | ***p* = 0.00038** |
|  | 20°C vs. 32°C | | ***p* = 1.5 x 10^-7^** |

*^1^* Different models with varying assumptions about treatments having different or sharing means and having different or sharing standard deviations (SD). *^2^* Compared to the winning model, more complex models have insignificant p-values (meaning the additional parameter(s) do not explain the data better) and less complex models have significant p-values. All p-values are compared to the winning model and are nested, log-likelihood ratio tests. Aikaike weights (calculated from ΔAIC values) estimate the likelihood of the model being best, given the suite of competing models.

**Table S6: PD (probability density) values for traits**. PD-values for host foraging rate (*f*), growth rate of parasites (*g_p_*), and transmission potential. PD-values are pairwise comparisons made using the 10,000 bootstrapped samples of one treatment and the best estimate of another. Accordingly, there are two values for each comparison (i.e., which treatment provides the bootstrapped distribution and which provides the best estimate). PD-values are analogous to *p*-values, but bounded between 0<PD<0.5. If PD=0, the best estimate is more extreme than all values in the bootstrapped distribution; if PD=0.5, the best estimate falls in the center of the distribution. We considered treatments significantly different (bold, below) if PD<0.025 (equivalent to a two-tailed test where α = 0.05).

| **Parameter** | | **Treatment Contrast**  **(best estimate vs. distribution)** | **PD-value** |
| --- | --- | --- | --- |
| Foraging rate (*f*) | | 20°C vs. 25°C | **PD = 0** |
| (Fig. 2B) | | 25°C vs. 20°C | **PD = 0** |
|  | | 25°C vs. 30°C | PD = 0.11 |
|  | | 30°C vs. 25°C | PD = 0.30 |
|  | | 20°C vs. 30°C | **PD = 0** |
|  | | 30°C vs. 20°C | **PD = 0** |
| Growth rate of parasites (*g_p_*) | | 20°C vs. 26°C | PD = 0.22 |
| (Fig 3B) | | 26°C vs. 20°C | PD = 0.15 |
|  | | 26°C vs. 32°C | PD = 0.21 |
|  | | 32°C vs. 26°C | PD = 0.33 |
|  | | 20°C vs. 32°C | PD = 0.52 |
|  | | 32°C vs. 20°C | PD = 0.50 |
| Transmission potential | | 20°C vs. 25/26°C | **PD = 0.0014** |
| (Fig. 5A) | | 25/26°C vs. 20°C | **PD = 0.017** |
|  | | 25/26°C vs. 30/32°C | **PD = 0.0001** |
|  | | 30/32°C vs. 25/26°C | **PD = 0** |
|  | | 20°C vs. 30/32°C | PD = 0.056 |
|  | | 30/32°C vs. 20°C | PD = 0.11 |

**Table S7: PD (probability density) values for transmission potential ‘sensitivity analysis’ calculations.** PD-values are pairwise comparisons made using the 10,000 bootstrapped samples of one treatment and the best estimate of another. Accordingly, there are two values for each comparison (i.e., which treatment provides the bootstrapped distribution and which provides the best estimate). PD-values are analogous to *p*-values, but bounded between 0<PD<0.5. If PD=0, the best estimate is more extreme than all values in the bootstrapped distribution; if PD=0.5, the best estimate falls in the center of the distribution. We considered treatments significantly different (bold, below) if PD<0.025 (equivalent to a two-tailed test where α = 0.05).

| **Parameter** | **Treatment Contrast**  **(point vs. CI distribution)** | **PD-value** |
| --- | --- | --- |
| Transmission potential without *f* | 20°C vs. 25/26°C | PD = 0.042 |
| (Fig. 5B) | 25/26°C vs. 20°C | PD = 0.12 |
|  | 25/26°C vs. 30/32°C | **PD = 0.0002** |
|  | 30/32°C vs. 25/26°C | **PD = 0** |
|  | 20°C vs. 30/32°C | **PD = 0.020** |
|  | 30/32°C vs. 20°C | **PD = 0.0075** |
| Transmission potential without *u* | 20°C vs. 25/26°C | **PD = 0.0025** |
| (Fig. 5C) | 25/26°C vs. 20°C | **PD = 0.013** |
|  | 25/26°C vs. 30/32°C | **PD = 0** |
|  | 30/32°C vs. 25/26°C | **PD = 0** |
|  | 20°C vs. 30/32°C | PD = 0.04 |
|  | 30/32°C vs. 20°C | PD = 0.118 |
| Transmission potential without *σ* | 20°C vs. 25/26°C | **PD = 0.0013** |
| (Fig. 5D) | 25/26°C vs. 20°C | **PD = 0.023** |
|  | 25/26°C vs. 30/32°C | **PD = 0.0003** |
|  | 30/32°C vs. 25/26°C | **PD = 0** |
|  | 20°C vs. 30/32°C | PD = 0.15 |
|  | 30/32°C vs. 20°C | PD = 0.095 |
| Transmission potential without *ρ* | 20°C vs. 25/26°C | **PD = 0.013** |
| (Fig. 5E) | 25/26°C vs. 20°C | PD = 0.15 |
|  | 25/26°C vs. 30/32°C | PD = 0.056 |
|  | 30/32°C vs. 25/26°C | **PD = 0.013** |
|  | 20°C vs. 30/32°C | PD = 0.50 |
|  | 30/32°C vs. 20°C | PD = 0.53 |
| Transmission potential without *φ* | 20°C vs. 25/26°C | **PD = 0.0003** |
| (Fig. 5F) | 25/26°C vs. 20°C | **PD = 0.0016** |
|  | 25/26°C vs. 30/32°C | **PD = 0.0006** |
|  | 30/32°C vs. 25/26°C | **PD = 0.0001** |
|  | 20°C vs. 30/32°C | PD = 0.39 |
|  | 30/32°C vs. 20°C | PD = 0.34 |
